# Putting bandits into context: How function learning supports decision making

**DOI:** 10.1101/081091

**Authors:** Eric Schulz, Emmanouil Konstantinidis, Maarten Speekenbrink

## Abstract

We introduce the contextual multi-armed bandit task as a framework to investigate learning and decision making in uncertain environments. In this novel paradigm, participants repeatedly choose between multiple options in order to maximise their rewards. The options are described by a number of contextual features which are predictive of the rewards through initially unknown functions. From their experience with choosing options and observing the consequences of their decisions, participants can learn about the functional relation between contexts and rewards and improve their decision strategy over time. In three experiments, we explore participants’ behaviour in such learning environments. We predict participants’ behaviour by context-blind (mean-tracking, Kalman filter) and contextual (Gaussian process and linear regression) learning approaches combined with different choice strategies. Participants are mostly able to learn about the context-reward functions and their behaviour is best described by a Gaussian process learning strategy which generalizes previous experience to similar instances. In a relatively simple task with binary features, they seem to combine this learning with a “probability of improvement” decision strategy which focuses on alternatives that are expected to lead to an improvement upon a current favourite option. In a task with continuous features that are linearly related to the rewards, participants seem to more explicitly balance exploration and exploitation. Finally, in a difficult learning environment where the relation between features and rewards is non-linear, some participants are again well-described by a Gaussian process learning strategy, whereas others revert to context-blind strategies.

## Introduction

Imagine you have recently arrived in a new town and need to decide where to dine tonight. You have visited a few restaurants in this town before and while you have a current favourite, you are convinced there must be a better restaurant out there. Should you revisit your current favourite again tonight, or go to a new one which might be better, but might also be worse? This is an example of the exploration exploitation dilemma (e.g., Cohen, McClure, & Yu, 2007; Laureiro-Martínez, Brusoni, & Zollo, 2010; Mehlhorn et al., 2015): should you exploit your current but incomplete knowledge to pick an option you think is best, or should you explore something new and improve upon your knowledge in order to make better decisions in the future? While exploration is risky, in this case it is not blind. Over the years, you have visited many restaurants and you know for instance that better restaurants generally have more customers, a good ambiance, and are not overly cheap. So you walk around town, noting of each restaurant you pass how busy it is, how nice it looks, the price of the items on the menu, etc. At the end of a long walk, you finally sit down in a restaurant; one you never visited before but predicted to be best based on numerous features such as neighbourhood, clientéle, price, and so forth.

The exploration-exploitation dilemma tends to be studied with so-called multi-armed bandit tasks, such as the Iowa gambling task (e.g., Bechara, Damasio, Tranel, & Damasio, 2005; Steyvers, Lee, & Wagenmakers, 2009). These are tasks in which people are faced with a number of options, each having an associated average reward. Initially, these average rewards are unknown and people can only learn about the reward of an option by choosing it. Through experience, people can learn which are the good options and use this knowledge in the attempt to accumulate as much reward as possible. However, as our restaurant example above shows, many real-life situations are richer than such simple multi-armed bandit tasks. Options tend to have numerous features (e.g., number of customers and menu prices in the restaurant example) which are predictive of their associated reward. With the addition of informative features, the decision problem can be termed a *contextual* multi-armed bandit (henceforth CMAB; Li, Chu, Langford, & Schapire, 2010). While these kinds of tasks are ubiquitous in daily life, they are rarely studied within the psychological literature. This is unfortunate, as CMAB tasks encompass two important areas of cognition: experience-based decision making (Barron & Erev, 2003; Hertwig & Erev, 2009; Speekenbrink & Konstantinidis, 2015) and function learning (DeLosh, Busemeyer, & McDaniel, 1997; Kalish, Lewandowsky, & Kruschke, 2004; Speekenbrink & Shanks, 2010). Both topics have been studied extensively (see e.g., Newell, Lagnado, & Shanks, 2015, for an overview), but commonly in isolation.

Learning and decision making within contextual multi-armed bandit tasks generally requires two things: learning a function that maps the observed features of options to their expected rewards, and a decision strategy that uses these expectations to choose between the options. Function learning in CMAB tasks is important because it allows one to generalize previous experiences to novel situations. For example, it allows one to predict the quality of a new restaurant from experiences with other restaurants with a similar number of customers and a similarly priced menu. The decision strategy is important because not only should you attempt to choose options that are currently most rewarding, but you should also take into account how much you can learn in order to make good choices in the future. In other words, you should take into account the exploration-exploitation trade-off, where exploration here means learning about the function that relates features to rewards.

In what follows, we will describe the contextual multi-armed bandit paradigm in more detail and propose several models to describe how people may solve CMAB tasks. We will then describe three experiments which explore how people perform within three variants of a CMAB task. We show that participants are able to learn within the CMAB, approximating the function in a close-to-rational way (Lucas, Griffiths, Williams, & Kalish, 2015; Srinivas, Krause, Kakade, & Seeger, 2009) and using their knowledge to sensitively balance exploration and exploitation. However, the extent to which participants are able to learn the underlying function crucially depends on the complexity of the task. In summary, we make the following contributions:

1. We introduce the contextual multi-armed bandit as a psychological paradigm combining both function learning and decision making.
2. We model and predict learning in CMABs using Gaussian processes regression, a powerful framework that generalizes important psychological models which were previously proposed to describe human function learning.
3. We show that participants sensibly choose between options according to their expectations (and attached uncertainty) while learning about the underlying functions.

## Contextual multi-armed bandits

A contextual multi-armed bandit task is a game in which on each round, an agent is presented with a context (a set of features) and a number of options which each offer an unknown reward. The expected rewards associated to each option depend on the context through an unknown function. The context can contain general features that apply to all options (e.g., the city the restaurants are in) or specific features that apply to single options (e.g., the exact menu and its price). The agent’s task is to choose those options that will accumulate the highest reward over all rounds of the game. The rewards are stochastic, such that even if the agent had complete knowledge of the task, a choice would still involve a kind of gamble. In this respect, choosing an option can be seen as choosing a slot machine (a one-armed bandit) to play, or, equivalently, choosing which arm of a multi-armed bandit to play. After choosing an option in a round, the agent receives the reward of the chosen option but is not informed of the foregone rewards that could have been obtained from the other options. For an agent who ignores the context, the task would appear as a restless bandit task (e.g., Speekenbrink & Konstantinidis, 2015), as the rewards associated with an arm will vary over time due to the changing context. However, learning the function that maps the context to (expected) rewards will make these changes in rewards predictable and thereby choosing the optimal arm easier. In order to choose wisely, the agent should thus learn about the underlying function. Sometimes, this may require her to choose an option which is not expected to give the highest reward on a particular round, but one that might provide useful information about the function, thus choosing to explore rather than to exploit.

Contextual multi-armed bandit tasks provide us with a scenario in which a participant has to learn a function in order to maximize the outputs of that function over time by making wise choices. They are a natural extension of both the classic multi-armed bandit task, which is a CMAB with an invariant context throughout, and the restless bandit task, which is a CMAB with time as the only contextual feature.

While the CMAB is novel in the psychological literature (though see Schulz, Konstantinidis, & Speekenbrink, 2015; Stojic, Analytis, & Speekenbrink, 2015), where few tasks explicitly combine function learning and experience-based decision making, there are certain similarities with tasks used in previous research. For example, recent studies in experience-based decision-making provided participants with descriptions about the underlying distributions that generate rewards (e.g., Lejarraga & Gonzalez, 2011; Weiss-Cohen, Konstantinidis, Speekenbrink, & Harvey, 2016). Just as in the CMAB, this presents a naturalistic decision environment in which different sources of information (e.g., descriptions and participants’ own experience) need to be integrated in order to choose between alternatives or courses of action.

Another related paradigm is multiple cue probability learning (MCPL, Kruschke & Johansen, 1999; Speekenbrink & Shanks, 2008) in which participants are shown an array of cues that are probabilistically related to an outcome and have to learn the underlying function mapping the cues’ features to expected outcomes. Especially when the outcome is a categorical variable, such as in the well-known “Weather Prediction Task” (Gluck, Shohamy, & Myers, 2002; Speekenbrink, Channon, & Shanks, 2008), making a prediction is structurally similar to a decision between multiple arms (possible predictions) that are rewarded (correct prediction) or not (incorrect prediction). Just as in the CMAB, multiple-cue probability learning and probabilistic category learning tasks require people to learn a function which maps multiple cues or features to expected outcomes. An important difference however is that in these latter tasks there is a strong dependency between the options: there is only one correct prediction, and hence there is a perfect (negative) correlation between the rewards for the options. Whether a current choice was rewarded or not thus provides information about whether the non-chosen options would have been rewarded. This dependency weakens the need for exploration, especially when the outcome is binary, in which case there is no need for exploration at all. In CMAB tasks, there is a stronger impetus for exploration, as the rewards associated to arms are generally conditionally independent, given the context. Knowing that a particular option was rewarded thus does not provide immediate information whether another option would have been rewarded. Another major difference is that MCPL tasks generally require participants to learn the whole function. In CMAB tasks, learning the function is only necessary insofar as it helps to make better decisions. To solve the exploration-exploitation dilemma, it may suffice to learn the function well only in those regions that promise to produce high rewards. Moreover, as we will see later, each option can be governed by its own function relating context to rewards. To our knowledge, simultaneous learning of mul-tiple functions has not previously been investigated.

Another area of related research comes from the associative learning literature, where it has been shown that context can act as an additional cue to maximize reward (cf Bouton & King, 1983; Gershman, Blei, & Niv, 2010). In one example of this, Gershman and Niv (2015) showed how generalization based on context (the average reward of options in an environment) can explain how participants react to novel options in the same environment, such that a high-reward context leads people to approach novel options, while a low-reward context leads to avoidance of novel options. The CMAB paradigm introduced here is related to such situations, but instead of a single, constant context, varies the contexts such that good performance requires learning the underlying contextual function.

## Models of learning and decision making

Formally, we can describe a CMAB as a game in which on each round *t* = 1,…, *T*, an agent observes a context 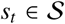 from the set 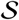 of possible contexts, and has to choose an arm 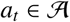 from the set 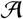 of all arms of the multi-armed bandit. After choosing an arm, the agent receives a reward

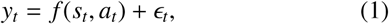

and it is her goal to choose those arms that will produce the highest accumulated reward

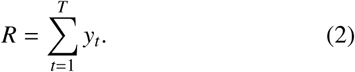

over all rounds. The function *f* is initially unknown and can only be inferred from the rewards received after choosing arms in the encountered contexts.

To perform well in a CMAB task, an agent needs to learn a model of the function *f* from experience, and on each round use this model to predict the outcomes of the available actions and choose the arm with the highest predicted outcome. We can thus distinguish between a learning component, formalized as a learning model which estimates the function relating rewards to contexts and actions, and a decision or acquisition component that uses the learned model to determine the best subsequent decisions. These work together as shown in Algorithm 1 (see also Brochu, Cora, & De Freitas, 2010).

### Algorithm 1 General CMAB-algorithm

A learning model 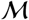 tries to learn the underlying function *f* by mapping the current expectations and their attached uncertainties to choices via an acquisition function acq.

**Require:** A model 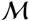 of the function *f*, an acquisition function acq, previous observations 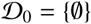

 **for** *t* = 1, 2,…, *T* **do**

   **Choose arm** 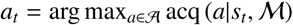

   **Observe reward** *y_t_* = *f* (*s_t_, a_t_*) + *ϵ_t_*

   **Update** Augment the data 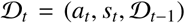 and update the model 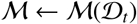

 **end for**

This formalization of an agent’s behaviour requires us to capture two things: (a) a representation or model 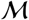 of the assumed underlying function that maps the given context to expected outcomes and (b) an acquisition function acq that evaluates the utility of choosing each arm based on those expected outcomes and their attached uncertainties. Here, the model defines the learning process and the acquisition function the way in which outputs of the learned model are mapped onto choices^1^. In the following, we will describe a number of instantiations of these two components.

### Models of learning

Technically, a function is a mapping from a set of input values to a set of output values, such that for each input value, there is a single output value (also called a many-to-one mapping as different inputs can provide the same output). Psychological research on how people learn such mappings has normally followed a paradigm in which participants are presented with input values and asked to predict the corresponding output value. After their prediction, participants are presented with the true output value, which is often corrupted by additional noise. Through this outcome feedback, people are thought to adjust their internal representation of the underlying function. In psychological theories of function learning, these internal representations are traditionally thought to be either *rule-based* or *similarity-based*. Rule-based theories (e.g., Carroll, 1963; Koh & Meyer, 1991) conjecture that people learn a function by assuming it belongs to an explicit parametric family, for example linear, polynomial, or power-law functions. Outcome feedback allows them to infer the parameters of the function (e.g., the intercept and slope of a linear function). This approach attributes a rich set of representations (parametric families) to learning agents, but tends to ignore how people choose from this set (how they determine which parametric family to use). Similarity-based theories (e.g., Busemeyer, Byun, Delosh, & McDaniel, 1997) conjecture that people learn a function by associating observed input values to their corresponding output values. When faced with a novel input value, they form a prediction by relying on the output values associated to input values that are similar to the novel input value. While this approach is domain general and does not require people to assume a parametric family a priori, similarity-based theories have trouble explaining how people readily generalize their knowledge to novel inputs that are highly dissimilar to those previously encountered.

Research has indicated that neither approach alone is sufficient to explain human function learning. Both approaches fail to account for the finding that some functional forms, such as linear ones, are much easier to learn than others, such as sinusoidal ones (McDaniel & Busemeyer, 2005). This points towards an initial bias towards linear functions, which can be overcome through sufficient experience. They also fail to adequately predict how people extrapolate their knowledge to novel inputs (DeLosh et al., 1997).

In order to overcome some of the aforementioned problems, hybrid versions of the two approaches have been put forward (McDaniel & Busemeyer, 2005). One such hybrid is the *extrapolation-association model* (EXAM, DeLosh et al., 1997), which assumes a similarity-based representation for interpolation, but simple linear rules for extrapolation. Although *EXAM* effectively captures the human bias towards linearity and accurately predicts human extrapolations over a variety of relationships, it cannot account for the human capacity to generate non-linear extrapolations (Bott & Heit, 2004). The *population of linear experts model* (POLE, Kalish et al., 2004) is set apart by its ability to capture knowledge partitioning effects; based on acquired knowledge, different functions can be learned for different parts of the input space. Beyond that, it demonstrates a similar ordering of error rates to those of human learners across different tasks (McDaniel, Dimperio, Griego, & Busemeyer, 2009). Recently, Lucas et al. (2015) proposed Gaussian process regression as a rational approach towards human function learning. Gaussian process regression is a Bayesian non-parametric model which unifies both rule-based and similarity-based theories of function learning. Instead of assuming one particular functional form, Gaussian process regression is based on a model with a potentially infinite number of parameters, but parsimoniously selects parameters through Bayesian inference. As shown by Lucas et al., a Gaussian process regression model accounts for many of the previous empirical findings on function learning. Following this approach, we will conceptualize function learning in a CMAB as Gaussian process regression. We contrast this with context-blind learning which tries to directly learn the expected reward of each option without taking the contextual features into account.

#### Contextual learning through Gaussian process regression

In the following, we will assume that the agents learns a separate function *f_j_*(*s*) that maps contexts *s* to rewards *y* for each option *j*. Gaussian process regression is a non-parametric Bayesian solution to function learning which starts with a prior distribution over possible functions and, based on observed inputs and outputs of the function, updates this to a posterior distribution over all functions. In Gaussian process regression, *p*( *f_j_*), the distribution over functions, is defined by a Gaussian process (GP). Technically, a GP is a stochastic process such that the marginal distribution of any finite collection of observations generated by it is a multivariate Gaussian distribution (see Rasmussen, 2006). A GP is parametrized by a mean function *m_j_*(*s*) and a co-variance function, also called kernel, *k_j_*(*s*, *s*′):

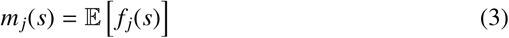

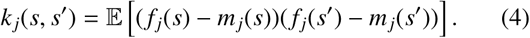

In the following, we will focus on the computations for a single option (and hence single function) and suppress the subscripts *j*. Suppose we have collected rewards **y***_t_* = [*y*_1_, *y*_2_,…, *y_t_*]^┬^ for arm *j* in contexts **s***_t_* = {*s*_1_,…, *s_t_*}, and we assume

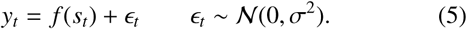

Given a GP prior on the functions

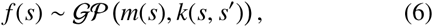

the posterior over *f* is also a GP:

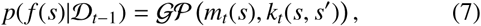

where 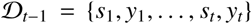 denotes the set of observations (contexts and rewards) of the function *f*. The posterior mean and kernel function are

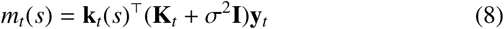

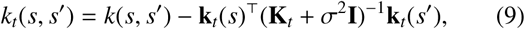

where **k***_t_*(*s*) = [*k*(*s*_1_, *s*),…, *k*(*s_t_*, *s*)]^┬^, **K***_t_* is the positive definite kernel matrix 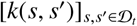, and **I** the identity matrix. Note that the posterior variance of *f* for context *s* can be computed as

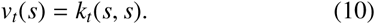

This posterior distribution can also be used to derive predictions about each arm’s rewards given the current context, that are also assumed to be normally distributed.

A key aspect of a GP model is the covariance or kernel function *k*. The choice of a kernel function corresponds to assumptions about the shape of the true underlying function. Among other aspects, the kernel determines the smoothness, periodicity, and linearity of the expected functions (c.f.Schulz, Tenenbaum, Duvenaud, Speekenbrink, & Gershman, 2016). Additionally, the choice of the kernel also determines the speed at which a GP model can learn over time (Schulz, Tenenbaum, Reshef, Speekenbrink, & Gershman, 2015). The kernel defines a similarity space over all possible contexts. As such, a GP can be seen as a similarity-based model of function learning, akin to exemplar models traditionally used to describe category learning (Nosofsky, 1986). However, by first mapping the contexts *s* via the kernel into a “feature space”, it is possible to rewrite the posterior mean of a GP as a linear combination of transformed feature values. From a psychological perspective, a GP model can in this way also be thought of as encoding “rules” mapping inputs to outputs. A GP can thus be simultaneously expressed as a similarity-based or rule-based model, thereby unifying the two dominant classes of function learning theories in cognitive science (for more details, see Lucas et al., 2015).

Different kernels correspond to different psychological assumptions about how people approach function learning. By choosing a *linear kernel*, the model corresponds directly to Bayesian linear regression. This kernel thus instantiates a relatively simple rule-based way of learning the underlying function, assuming it has a particular parametric shape, namely a linear combination of the contextual features. The *radial basis function kernel* (RBF, sometimes also called square(d) exponential or Gaussian kernel) postulates smooth but otherwise relatively unconstrained functions and is probably the most frequently used kernel in the Gaussian process literature. The RBF kernel contains a free parameter *λ*, referred to as the length scale, which determines the extent to which increasing the distance between two points reduces their correlation. The mathematical details of the two contextual models, corresponding to these two choices of kernel, as well as an illustration of the way in which they learn (i.e. update their prior distribution to a posterior distribution) are provided in Table 1.

**Table 1.**
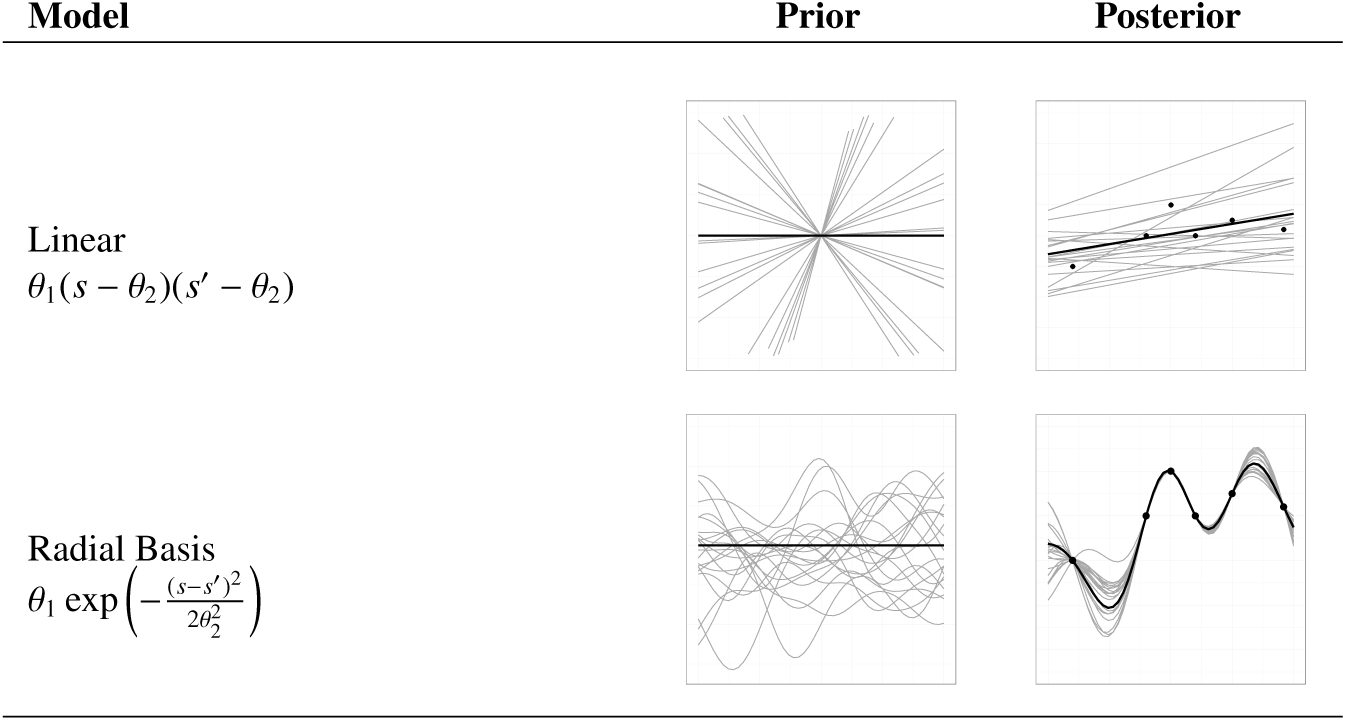
Details of the two contextual models used to model participants’ learning. Mathematical details of each model are provided in the “Model” column. For each model, prior samples of functions for a one-dimensional input are shown in the “Prior” column. The “Posterior” column shows posterior samples of the functions after the same set of 6 observations (dots).

| Model | Prior | Posterior |
| --- | --- | --- |
| Linear<br>$\theta_1(s - \theta_2)(s' - \theta_2)$ | | |
| Radial Basis<br>$\theta_1 \exp\left(-\frac{(s-s')^2}{2\theta_2^2}\right)$ | | |

#### Context-blind learning

To assess the extent to which people take the context into account, we contrast the contextual learning models above with two context-blind learning models that ignore the features and focus on the average reward of each option over all contexts.

The *Bayesian mean-tracking* model assumes that the average reward associated to each option is constant over time and simply computes a posterior distribution over the mean *μ_j_* of each option *j*. Here, we will implement a relatively simple version of such a model which assumes rewards are normally distributed with a known variance but unknown mean and the prior distribution for that mean is again a normal distribution. This implies that the posterior distribution for each mean is also a normal distribution:

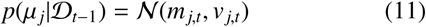

Here, the mean *m_j,t_* represents the currently expected outcome for a particular arm *j* and the variance *v_j,t_* represents the uncertainty attached to that expectation. The posterior distribution can be computed through a mean-stable version of the Kalman filter, which we will describe next.

Unlike the Bayesian mean tracking model, which computes the posterior distribution of a time-invariant mean *μ_j_* after each new observation, the Kalman filter is a suitable model for tracking a time-varying mean *μ_j,t_* which we here assume varies according to a simple random walk

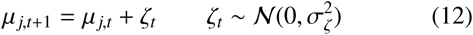

Such a Kalman filter model has been used to successfully describe participants’ choices in a restless bandit task (Speekenbrink & Konstantinidis, 2015) and has also been proposed as a model unifying many findings within the literature of context-free associative learning (Gershman, 2015; Kruschke, 2008). In this model, the posterior distribution of the mean is again a normal distribution

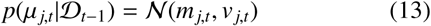

with mean

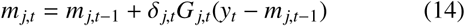

where *y_t_* is the received reward on trial *t* and *δ_j,t_* = 1 if arm *j* was chosen on trial *t*, and 0 otherwise. The “Kalman gain” term is computed as

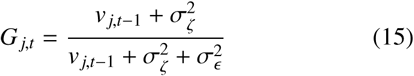

where *v_k,t_*, is the variance of the posterior distribution of the mean *μ_j,t_* is computed as

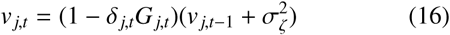

Prior means and variances were initialized to *m_j_*_,0_ = 0 and*v_j_*_,0_ = 1000, while the innovation variance 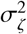 and errorvari-ance 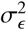 error free parameters. The Bayesian mean-tracking model is obtained from the Kalman filter model by setting the innovation variance to 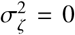, implying the underlyingmean is not assumed to change over time.

### Decision strategies

The aforementioned learning models each generate a predictive distribution, reflecting the rewards expected from choosing options in the current context. To model participants’ choices, we need a decision strategy that defines the current predictive means and variances are used to choose between options. In the psychological literature, popular decision rules that map current expectations onto choices are the softmax and *ϵ*-greedy rule (Sutton & Barto, 1998). These are rules which are based on a single expectation for each option. In the softmax rule, the probability of choosing an option is roughly proportional to the current expectations, while the *ϵ*-greedy rule chooses the maximum-expectancy option with probability 1 − *ϵ* and otherwise chooses with equal probability between the remaining options. Frequently, these rules ignore the uncertainty about the formed expectations, while rationally, uncertainty should guide exploration. Here, we follow Speekenbrink and Konstantinidis (2015) and define a broader set of decision rules that explicitly model how participants trade off between expectations and uncertainty. We will consider 4 different strategies to make decisions in a CMAB task based on the predictive distributions derived from the above learning models. The mathematical details of these are given in Table 2.

**Table 2.**
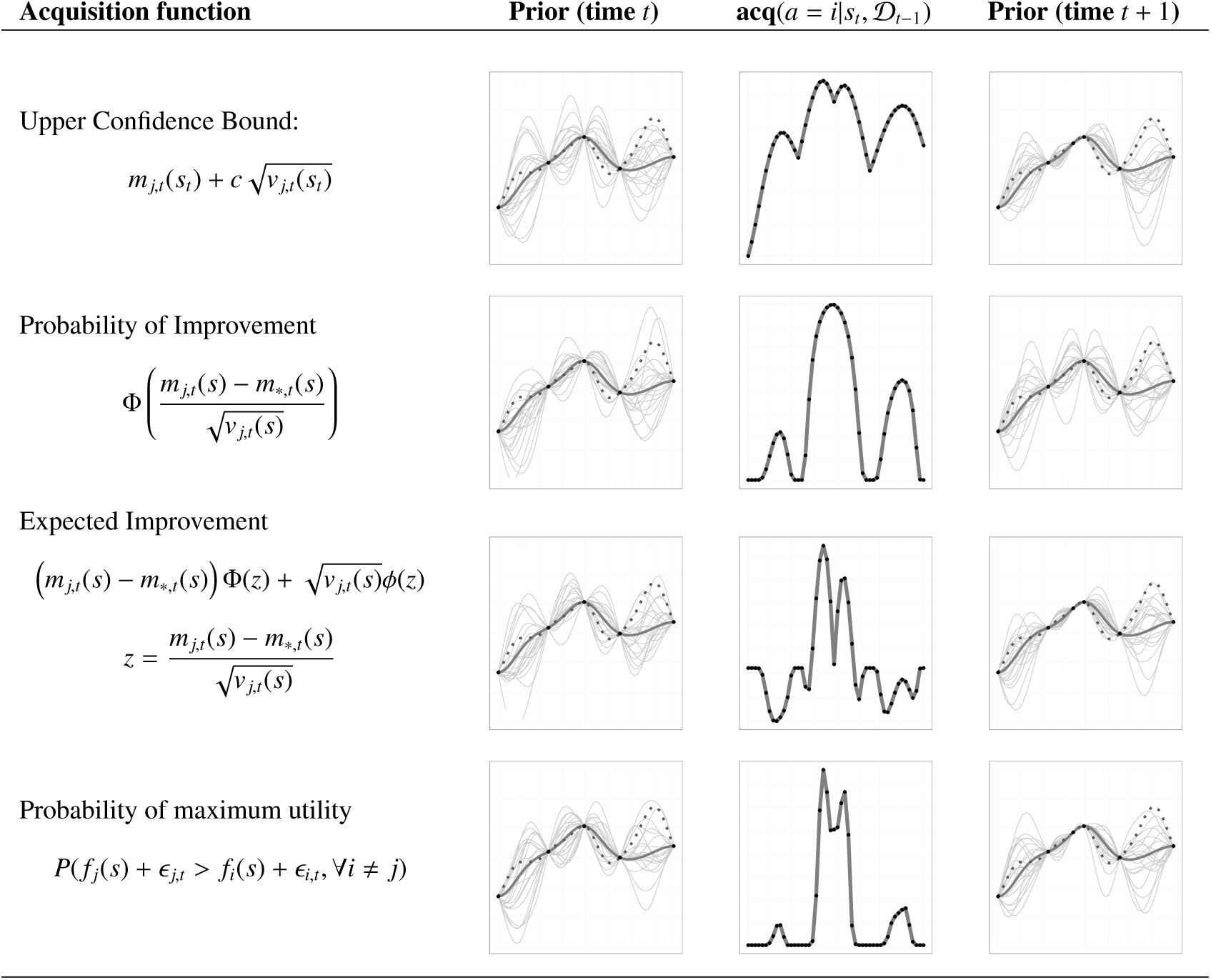
Acquisition functions used to model participants’ choices. Mathematical details are provided in the column “Acquisition function”. Here, m_j,t_(s) denotes the posterior mean of the function for context s and action j, and action j = * denotes the action currently believed to be optimal. Examples are provided for a problem where each action corresponds to choosing a one-dimensional input, after which the associated output can be observed. Prior samples from a Radial Basis kernel are shown in the “Prior (time t)” column. The utility of each potential action according to each acquisition function is shown in the “acq()” column. After choosing the action with the highest utility and observing the corresponding output, the Gaussian process is updated and used as a prior at the next time. Samples from this posterior are shown in the final column(“Prior (time t + 1)”).

The *upper confidence bound* (UCB) algorithm defines a trade-off between an option’s expected value and the associated uncertainty and chooses the option for which the upchoice:per confidence bound of the mean is highest. The UCB rule has been shown to perform well in many real world tasks(Krause & Ong, 2011). It has a free parameter *β*, which determines the width of confidence interval (for example, *β* = 1.96 would result in a 95% credible set). The UCB-algorithm can be described as a selection strategy with an exploration bonus, where the bonus dynamically depends on the confidence interval of the estimated mean reward at each time point. It is sometimes also referred to as optimistic sampling as it can be interpreted to inflate expectations with respect to the upper confidence bounds (Srinivas et al., 2009).

Another decision strategy is the *probability of improvement* (PoI) rule, which calculates the probability for each option to lead to an outcome higher than the option that is currently believed to have the highest expected value (Kushner, 1964). Intuitively, this algorithm estimates the probability of one option to generate a higher utility than another option and has recently been used in experiments involving multi-attribute choices (Gershman, Malmaud, Tenenbaum, & Gershman, 2016).

The PoI rule focusses solely on the probability that an option provides a higher outcome than another; whether the difference in outcomes is large or small does not matter. The *expected improvement* (EI) rule is similar to the PoI rule, but does take the magnitude of the difference in outcomes into account and compares options to the current favourite in terms of the expected increase of outcomes (Mockus, Tiesis, & Zilinskas, 1978).

The fourth decision strategy we consider is the *probability of maximum utility* (PMU) rule (Speekenbrink & Konstantinidis, 2015). This strategy chooses each option according to the probability that it results in the highest reward out of all options in a particular context. It can be seen as a form of probability matching (Neimark & Shuford, 1959) and can be implemented by sampling from each option’s predictive distribution once, and then choosing the option with the highest sampled pay-off. Even though this acquisition function seems relatively simple, it describes human choices in restless bandit tasks well (Speekenbrink & Konstantinidis, 2015). It is also closely related to Thompson sampling (May, Korda, Lee, & Leslie, 2012), which samples from the posterior distribution of the mean rather than the predictive distribution of rewards. Thus, while Thompson sampling “probability matches” the expected rewards of each arm, the probability of maximum utility rule matches to actual rewards that might be obtained^2^.

The first three decision rules (but not the PMU rule) are deterministic, while participants’ decisions are expected to be more noisy reflections of the decision rule. We therefore used a softmax transformation to map the value of each option according to the decision rule into probabilities of choice:

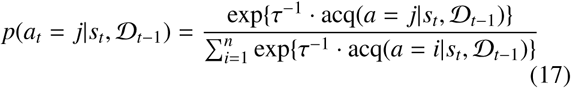

The temperature parameter *τ* > 0 governs how consistently participants choose according to the values generated by the different kernel-acquisition function combinations. As *τ* → 0 the highest-value option is chosen with a probability of 1 (i.e., arg max), and when *τ* → ∞, all options are equally likely, with predictions converging to random choice. We use *τ* as a free parameter, where lower estimates can be interpreted as more precise predictions about choice behaviour.

## General CMAB task

In our implementation of the CMAB task, participants are told they have to mine for “Emeralds” on different planets. Moreover, it is explained that at each time of mining the galaxy is described by 3 different environmental factors, “Mercury”, “Krypton”, and “Nobelium”, that have different effects on different planets. Participants are then told that they have to maximize their production of Emeralds over time by learning how the different environmental factors influence the planets and choosing the planet they think will produce the highest outcome in light of the available factors. Participants were explicitly told that different planets can react differently to specific environmental factors. A screen-shot of the CMAB task can be seen in Figure 1.

**Figure 1.**
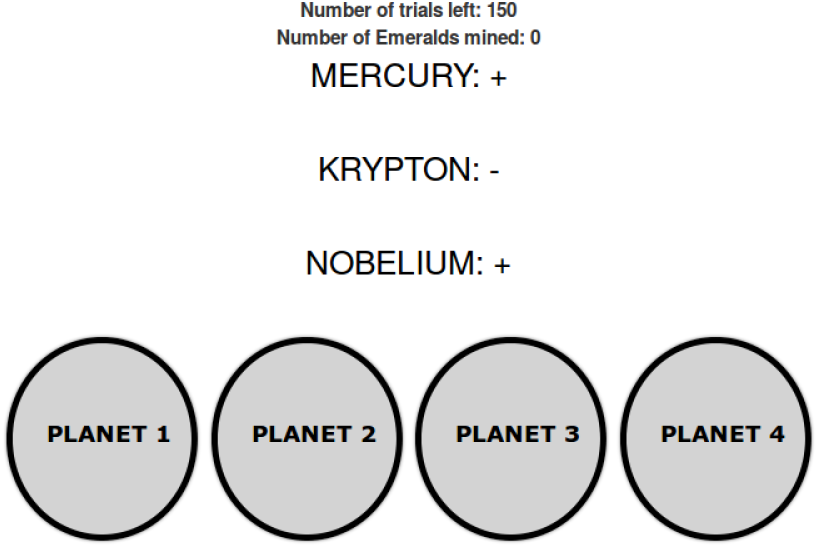
Screenshot of the CMAB task in Experiment 1.

As each planet responds differently to the contexts, they can be seen as arms of a multi-armed bandit that are related to the context by different functions. The reward of an option *j* is given as

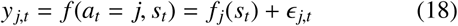

with 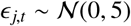. The task consists of 150 trials in which a random context is drawn and participants choose a planet to mine on^3^.

The three experiments differ in the functions *f_j_* and whether the environmental factors defining the context were binary or continuous. This is specified in more detail when describing the experiments. Source code for the experimental set-up is available online.^4^

## Model comparison

All models were compared in terms of their out-of-sample predictive performance, assessing the accuracy of their one-step-ahead predictions and comparing it to that of a *random model* which picks each option with the same probability. Our procedure is as follows: for each participant, we first fitted a given model by maximum likelihood to the first *t* − 1 trials with a differential evolution optimization algorithm (using 100 epochs, cf. Mullen, Ardia, Gil, Windover, & Cline, 2009). We then used this fitted model to predict the choice on trial *t*. As repeating this procedure for every trial is computationally expensive, we assess the models’ predictive accuracy for every participant on trials *t* = {10, 30, 50, 70, 90, 110, 130, 150}. The one-step-ahead predictive accuracy measure compares each model 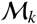 to a random model 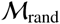:

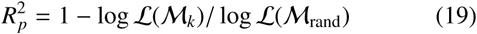

where 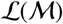 denotes the likelihood of model 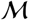 (i.e., the probability of a participants’ choices as predicted by fitted model 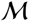). This measure is similar to McFadden’s pseudo-*R*^2^ (McFadden, 1973), although it uses the completely random model 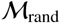 as comparison model, instead of the intercept-only regression model used in McFadden’s pseudo-*R*^2^. Just like McFadden’s measure, ours has values between 0 (accuracy equal to the random model) and 1 (accuracy infinitely larger than the random model).

## Experiment 1: CMAB with binary cues

The goal of the first experiment was to test whether participants can learn to make good decisions in a CMAB task. For this purpose, we set up a relatively simple contextual bandit scenario in which the contexts consist of binary features.

### Participants

Forty-seven participants (26 male) with an average age of 31.9 years (*S D* = 8.2) were recruited via Amazon Mechanical Turk and received $0.3 plus a performance-dependent bonus. The experiment took 12 minutes to complete on average and the average reward was $0.73±0.07.

### Task

There were four different arms that could be played (planets that could be mined). In addition, three discrete variables, *s_i,t_*, *i* = 1, 2, 3, were introduced as the general context. The three variables defining the contexts could either be on (*s_i,t_* = 1) or off (*s_i,t_* = −1). The outcomes of the four arms were dependent on the context as follows:

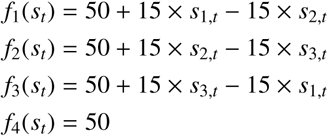

The assignment of these functions to the planets, and the order of the planets on screen, was the same for each participant.^5^

On each trial, the probability that a contextual feature was on or off was set to *p*(*s_i,t_* = 1) = *p*(*s_i,t_* = −1) = 0.5, making each of the 8 possible contexts equally likely to occur on a given trial. The functions *f_j_* were designed such that the expected reward of each arm over all possible contexts equals 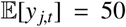. This means that the only way to gain higher rewards than the average of 50 is by learning how the contextual features influence the rewards. More formally, this implies that no arm achieves first-order stochastic dominance. Moreover, including the context-independent fourth arm that returns the mean with added noise helps us to distinguish even further between learning and not learning the context: this arm has the same expected value as all the other arms but a lower variance and therefore achieves second-order stochastic dominance over the other arms. As such, a context-blind and risk-averse learner would prefer this arm over time.

### Procedure

After giving their informed consent, participants received instructions to the experiment. Participants were told that they had to mine for “Emeralds” on different planets. Moreover, it was explained that at each time each of the 3 different environmental factors could either be on (**+**) or off (**-**) and had different effects on different planets. Participants were told that they had to maximize the overall production of Emeralds over time by learning how the different elements influence the planets and then picking the planet they thought would produce the highest outcome, given the status (on or off) of the elements. It was explicitly noted that different planets can react differently to different elements. After reading the instructions, participants performed the CMAB task. There were a total number of 150 trials and participants were paid $0.3 + total score/(150 × 100).

### Results

For all of the following analyses we report both frequentist and Bayesian test results. The latter are reported as Bayes factors, where *BF*_10_ quantifies the posterior probability ratio of the alternative hypothesis as compared to the null hypothesis (see Morey, Rouder, Jamil, & Morey, 2015). Unless stated otherwise, we use a Bayesian t-test (Morey & Rouder, 2011; Rouder, Speckman, Sun, Morey, & Iverson, 2009), with a Jeffreys-Zellner-Siow (JZS) prior with scale 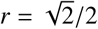.

#### Behavioural results

Participants gained 66.78 points (SD=13.02) per round on average throughout the task. Participants’ average scores were significantly above the chance level of 50 (*t*(46) = 8.83, *p* < 0.01). 34 out of 47 participants performed better than chance according to a simple t-test with *α* = 0.05 and *μ*_0_ = 50. Using a Bayesian meta-analytical t-test^6^ over all participants’ scores, we found a Bayes factor of *BF*_10_ = 68.34 indicating that the alternative hypothesis of participants performing better than chance was around 68 times more likely than chance performance. As such, participants seemingly learned to take the context into account, obtaining higher rewards than expected if they were ignoring the context.

Over time, participants made increasingly better choices (see Figure 2a), as indicated by a significant correlation between the average score (over participants) and trial number, *r* = 0.74, *p* < 0.01. Using a Bayesian test for correlations (Wetzels & Wagenmakers, 2012), we found a Bayes factor of *BF*_10_ = 6.01 when comparing the correlation to a mean of 0.27 out 47 participants had a significantly positive correlation between trial numbers and score at *α* = 0.05.

**Figure 2.**
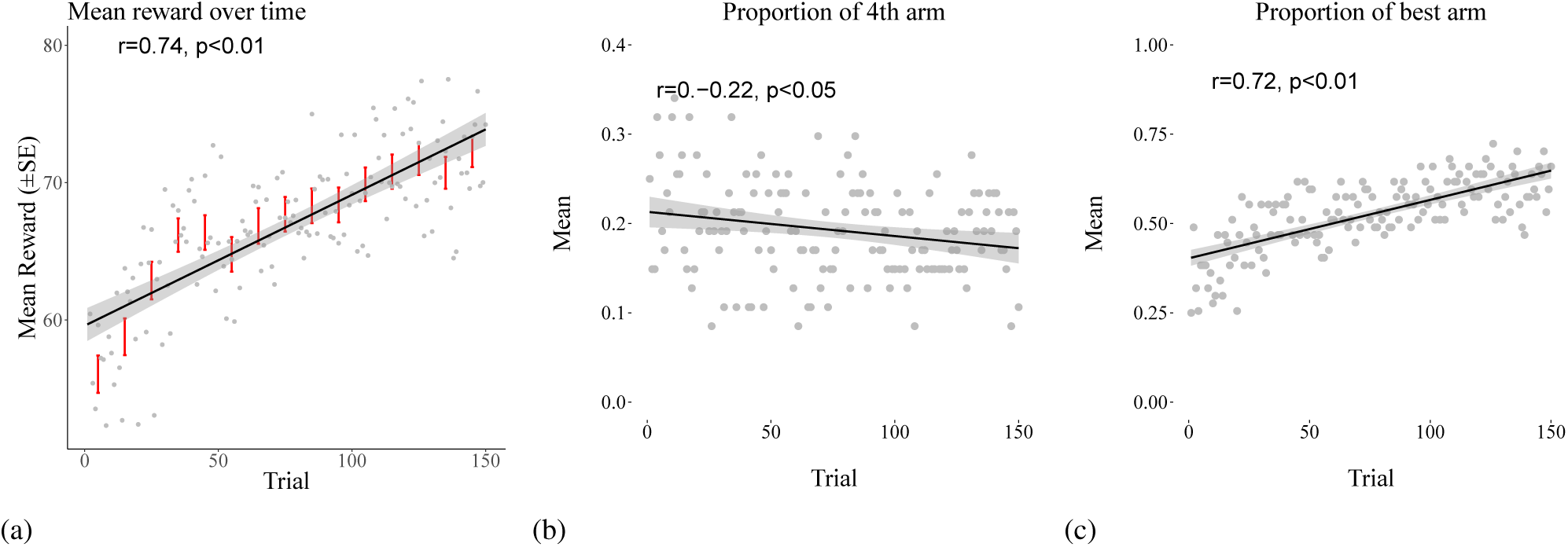
Results of the the continuous-linear CMAB task of Experiment 1. (a) average mean score per round, (b) proportion of choices of the 4th arm, and (c) proportion of choices of the best arm. Red error bars indicate standard error aggregated over 5 trials. Regression line is based on a least square regression including a 95% confidence level interval of the prediction line.

The proportion of participants choosing the non-contextual option (the option that did not respond to any of the contextual features, indicated as the 4th arm) decreased over time (*r* = −0.22, *p* < 0.05, *BF*_10_ = 58.8, Figure 2b), another indicator that participants learned the underlying functions. Finally, the proportion of participants choosing the best option for the current context increased during the task (*r* = 0.72, *p* < 0.01, *BF*_10_ = 263.2, see Figure 2a). Moreover, when assessing whether either outcomes or chosen arms on a trial *t* − 1 were predictive for a chosen arm on trial *t* in a hierarchical multinomial regression (where trials were nested within participants) with chosen arms as dependent variable, we found no significant relationship, again indicating that participants seemed to indeed learn the underlying function instead of using more simplistic (and in this case not very useful) heuristic memorization techniques such as testing similar arms in sequences or changing to a particular arm after a particular score.

#### Modelling results

To determine which combination of learning model and acquisition function best captures participants’ choices, we focus on one-step-ahead predictive comparisons. For each participant and model, we computed our pseudo-*R*^2^ at the eight test trials. Higher *R*^2^-values indicate better model performance. The results are shown in Figure 3.

**Figure 3.**
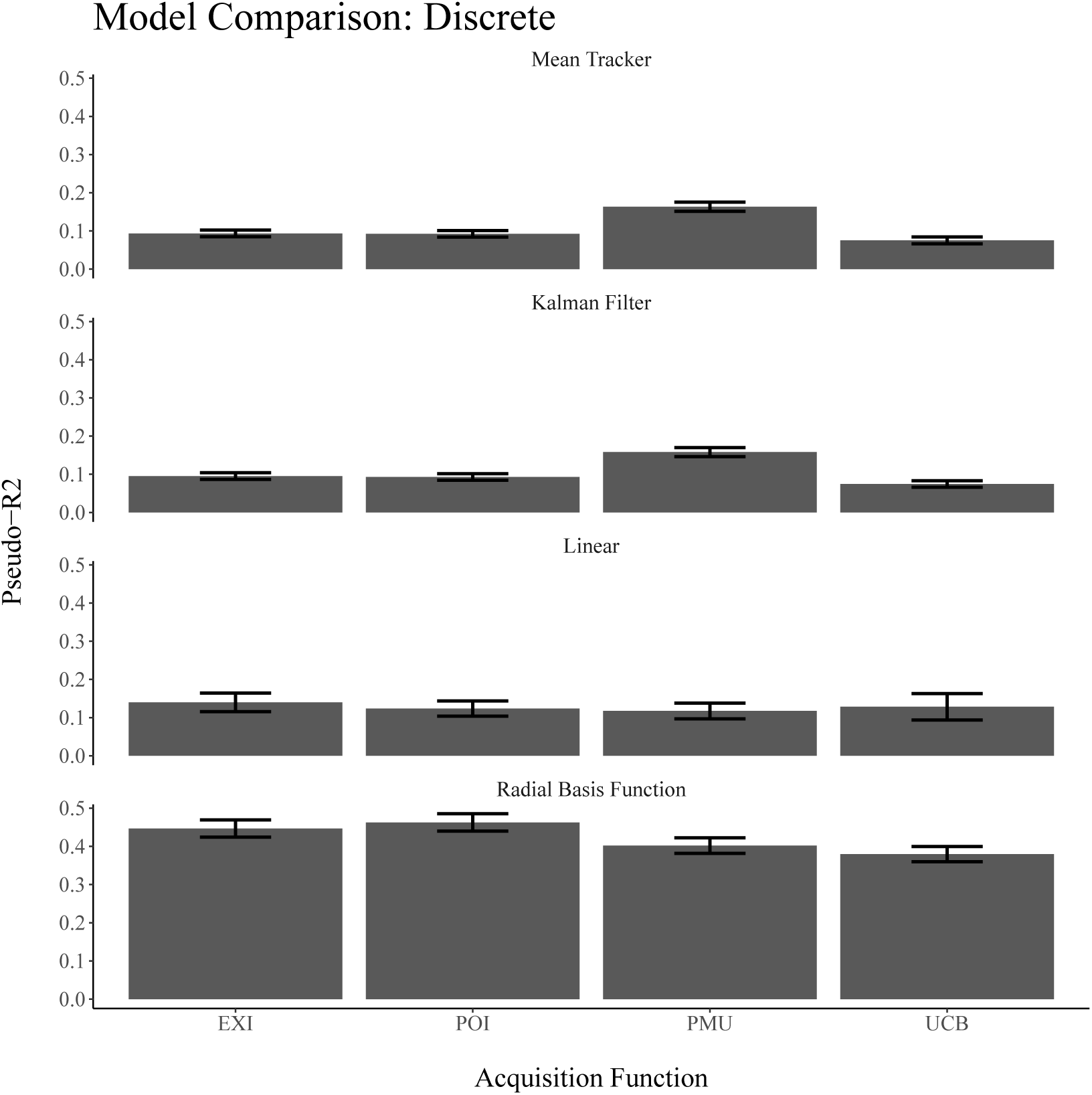
Predictive accuracy of the models for the CMAB task with discrete cues in Experiment 1. Error bars represent the standard error of the mean.

Overall, the best performing model was the GP learning model with a RBF kernel and the PoI decision rule. Aggregating over acquisition functions, the contextual models produced significantly better one-step-ahead predictions than the context-blind models (*t*(186) = 6.13, *p* < 0.01, *BF*_10_ = 1.9 × 10^4^). Additionally, the GP-model with an RBF kernel performed better than the linear model (*t*(92) = 7.23, *p* < 0.01, *BF*_10_ = 2.6 × 10^4^). Distinguishing the different acquisition functions turned out to be harder than comparing the different learning approaches. Aggregating over learning models, the probability of maximum utility strategy performed marginally better than all other acquisition functions (*t*(186) = 1.97, *p* < 0.05, *BF*_10_ = 2.3). Even though the probability of improvement acquisition function numerically predicted participants’ choices best out of all the acquisition functions when combined with the RBF kernel GP, this difference was not high (*t*(186) = 1.15, *p* > 0.05, *BF*_10_ = 0.24).

The median parameter estimates of the GP model over all acquisition functions per participant were extracted and are shown in Figure 4.

**Figure 4.**
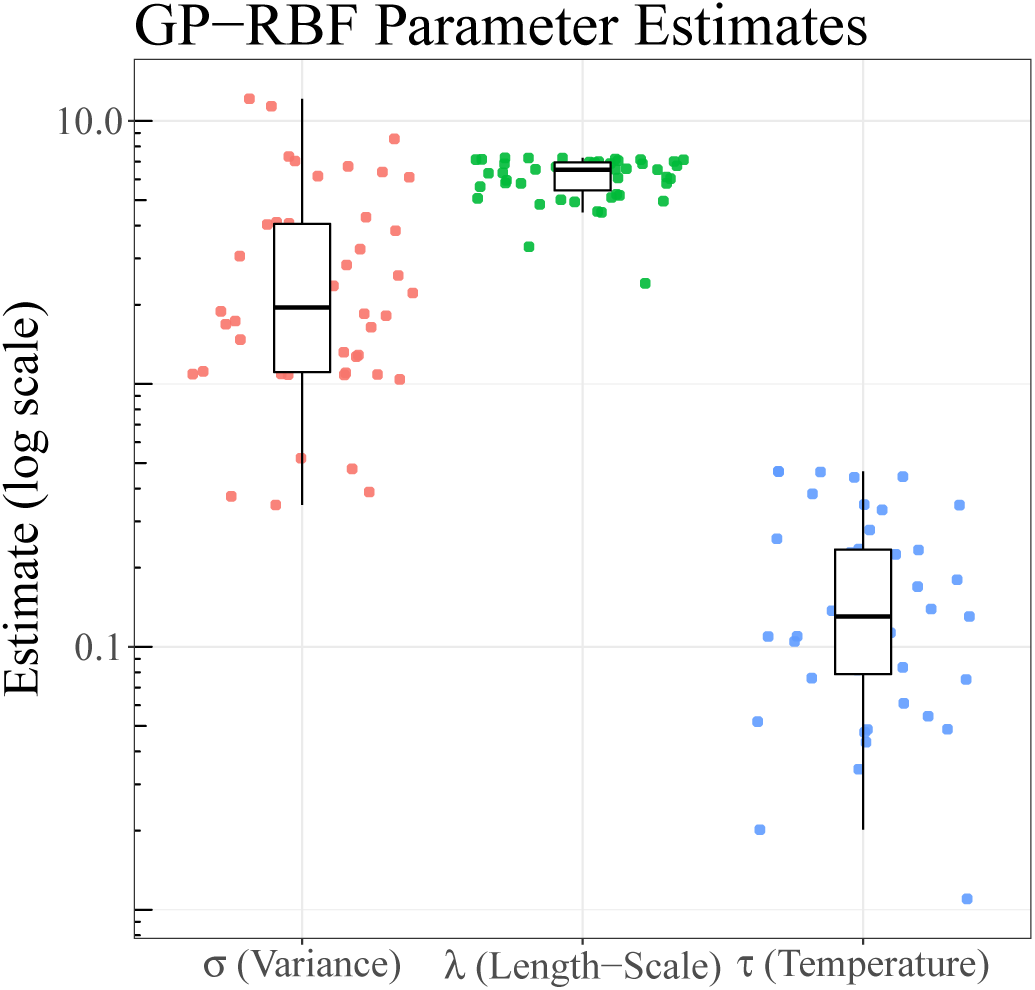
Parameter estimates of the error variance *σ*, the length-scale *λ*, and the temperature parameter *τ* for the GP-RBF model in Experiment 1. Dots show median parameter estimates per participant and boxplots show the median and inter-quartile range.

The median noise variance 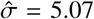 was reasonably close to the underlying observation noise variance of *σ* = 5, albeit smaller in general (*t*(46) = −4.7, *p* < 0.01, *BF*_10_ = 913.05); thus, participants seemed to underestimate the overall noise in the observed outcomes. The estimates of the length-scale parameter clustered around the mean value of 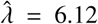. An RBF kernel can emulate a linear kernel by setting a very high length-scale. As the true underlying functions were linear in the experiment, we could thus expect high values for 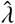. In that light, a value of six for the estimated length-scale seems surprisingly small, as it indicates that the dependencies between input points are expected to decay rather quickly, i.e. that participants generalized more locally than what was necessary. The overall temperature parameter was relatively low (mean estimate: 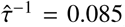), indicating that participants quite consistently chose the options with the highest predicted rewards.

According to the best fitting model in our cognitive modelling exercise, people learn the relation between context and outcomes by relying on a more general function approximator than just a linear regression (implemented as a linear kernel). By using a Probability of Improvement decision strategy, participants compare the option which is thought to have the highest average rewards in the current context, to relatively lesser known options in that context, determining how probable these are to provide a higher reward. This strategy is in agreement with prior findings in simpler multi-attribute choice tasks (for example, Carroll & De Soete, 1991).

## Experiment 2: Continuous-Linear CMAB

Experiment 1 contained only 8 unique contexts. This makes a memorization strategy feasible: participants may have simply memorized the expected rewards for each option in each context, rather than inferring a more general model of the underlying function. The goal of the second experiment was to assess whether the findings from Experiment 1 generalize to a task with a larger number of unique contexts, in which memorization of input-output pairs is less plausible. For this purpose, Experiment 2 used the same task as Experiment 1, but with continuous rather than binary features comprising the contexts.

### Participants

Fifty-nine participants (30 male) with a mean age of 32.4 (SD=7.8) were recruited via Amazon Mechanical Turk and received $0.3 as a basic reward and a performance-dependent bonus of up to $0.5. The experiment took 13 minutes on average to complete and the average reward was $0.69 ± 0.08.

### Task and Procedure

The task was identical to that of Experiment 1, only this time the context contained continuous features with an underlying linear function mapping inputs to outputs:

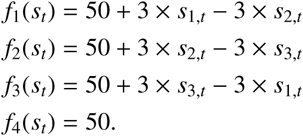

The values of the context variables *s_j,t_* were sampled randomly from a uniform distribution 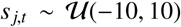. The values were rounded to whole numbers and shown in their numerical form to participants. As in the task of Experiment 1, the expected value (over all contexts) for each option was 50, so no option achieved first-order stochastic dominance, while the fourth option achieved second-order stochastic dominance as the variance of its rewards was the lowest.

### Results

#### Behavioral results

On average, participants earned 59.84 (SD = 9.41) points during the entire game, which is significantly higher than chance, *t*(58) = 7.17, *p* < 0.01. A hierarchical Bayesian t-test revealed that the alternative hypothesis of performing better than chance was *BF*_10_ = 53.88 more likely than the null hypothesis of chance performance. 29 participants performed better than chance overall as measure by individual t-tests with *α* = 0.05. Thus, as in Experiment 1, participants were able to take the context into account in order to increase their performance above chance level.

Performance increased over trials, *r* = 0.39, *t*(58) = 3.64, *p* < 0.01, although this was not as pronounced as in Ex-periment 1 (see Figure 5a). A hierarchical Bayesian t-test showed that participants’ correlations between score and trial number were *BF*_10_ = 15.44 more likely to be greater than 0 than lesser than or equal to 0, thus showing strong evidence for improvement over time. The correlation between trial number and score was significantly positive for 20 out of 59 participants.

**Figure 5.**
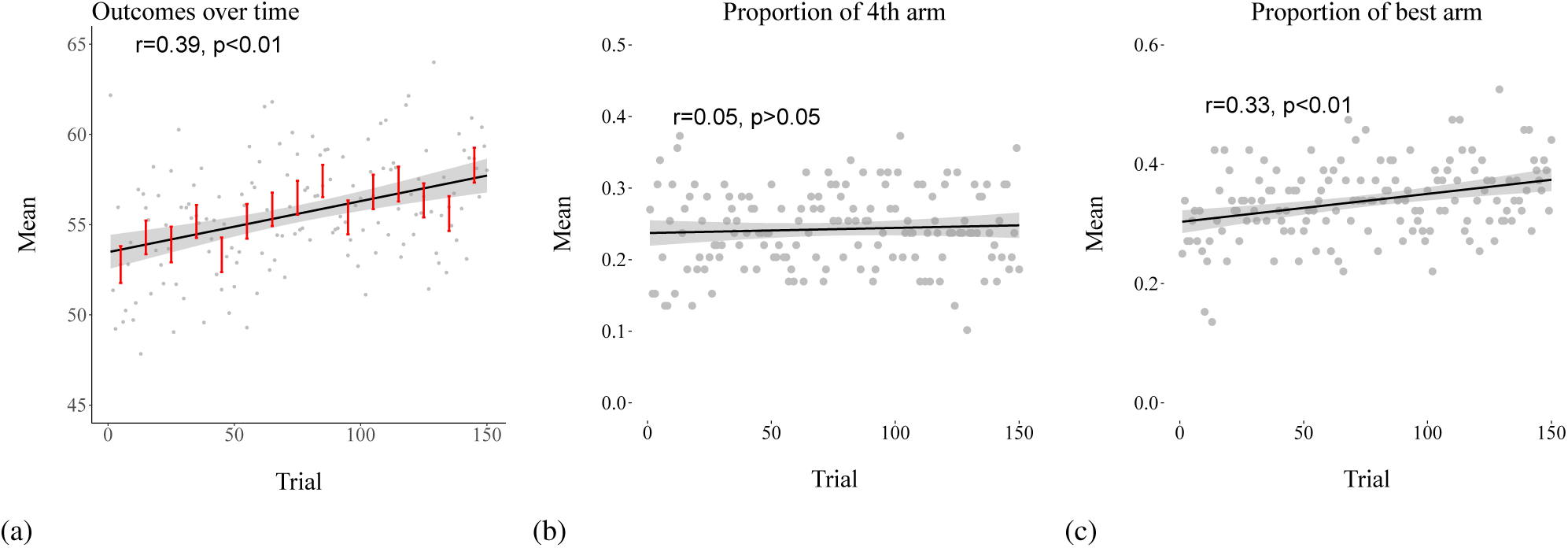
Results of the the continuous-linear CMAB task of Experiment 2. (a) average mean score per round, (b) proportion of choices of the 4th arm, and (c) proportion of choices of the best arm. Red error bars indicate standard error aggregated over 5 trials. Regression line is based on a least square regression including a 95% confidence level interval of the prediction line.

While the proportion of participants choosing the fourth option did not decrease significantly over time (*r* = 0.05, *p* > 0.05, *BF*_10_ = 0.01), the proportion of choosing the best option in the context did increase significantly over trials (*r* = 0.33, *p* < 0.01, *BF*_10_ = 18.87 see Figure 5c).

A hierarchical multinomial regression showed that neither the previously chosen arm nor the previously received reward was predictive of current choice (all *p* > 0.05). Thus, participants did not seem to rely on simply repeating choices or other more simple heuristics to determine their decisions.

#### Modelling results

Cross validation results are shown in Figure 6. The best performing model incorporates again a GP-RBF learning component, but now coupled with a UCB decision strategy. In this experiment, the contextual models did not significantly outperform the context-blind models (*t*(234) = −2.59, *p* < 0.01, *BF*_10_ = 0.12). However, this was mostly due to the linear model performing significantly worse than all the other learning models (*t*(234) = 2.37, *p* < 0.05, *BF*_10_ = 8.79). The GP-RBF model significantly outperformed all the other candidate learning models (*t*(234) = 5.63, *p* < 0.01, *BF*_10_ = 6.73). Thus, as in Experiment 1, participants were best predicted by a Gaussian Process learning model with a radial basis function kernel.

**Figure 6.**
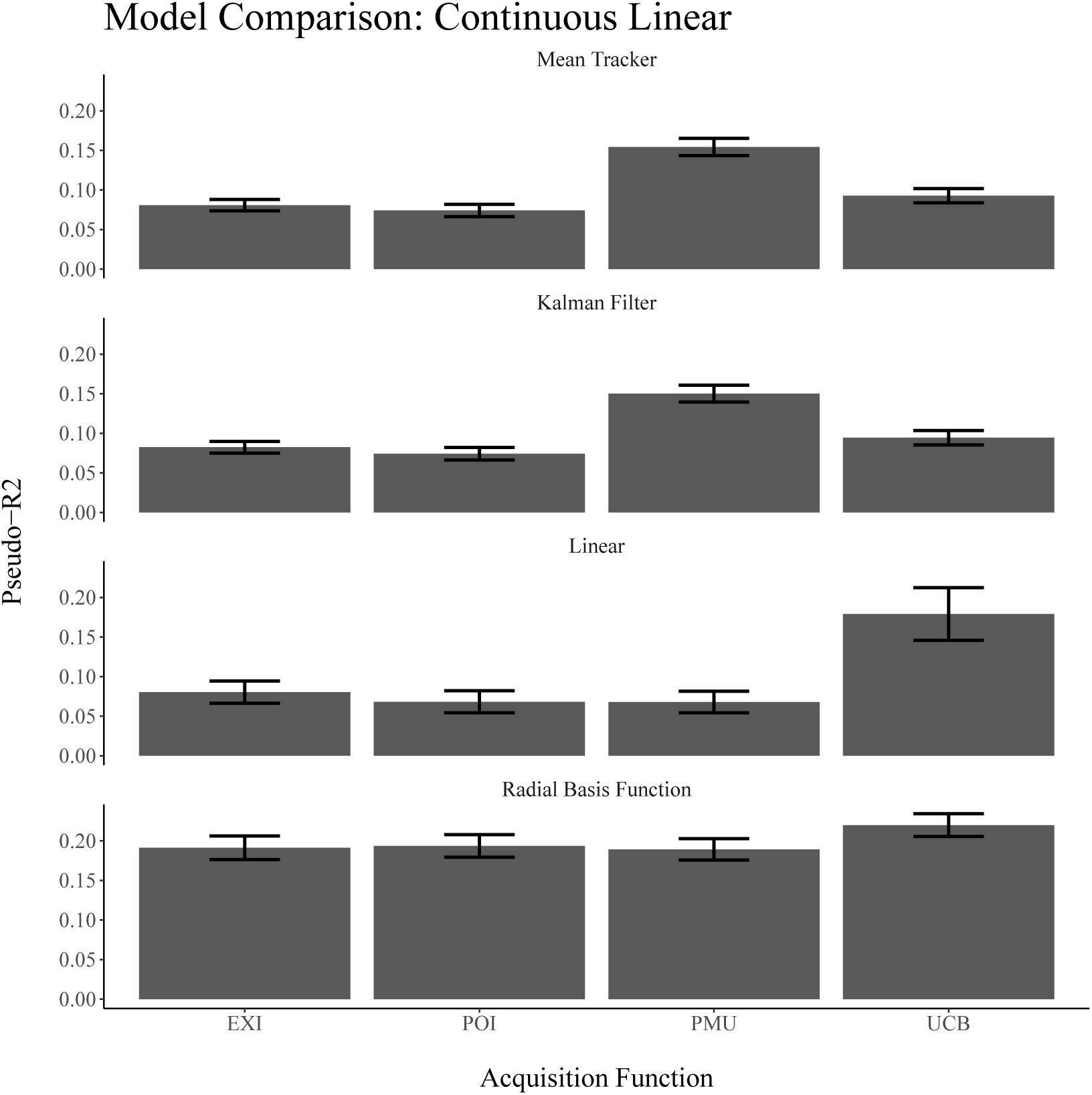
Predictive accuracy of the models for the CMAB task with continuous-linear cues in Experiment 2. Error bars represent the standard error of the mean.

The best performing decision strategy differs between the contextual and context-free models. The UCB strategy performed better than the other decision strategies for the contextual models, significantly so for the linear learning model, *t*(609) = 3.94, *p* < 0.01, *BF*_10_ = 7.45, but not signifi-cantly for the RBF-learning model, *t*(609) = 0.4, *p* > 0.05, *BF*_10_ = 3.62. For the context-free learning models, the probability of maximum utility acquisition function provided the best predictive performance for both the Bayesian mean tracker (*t*(614) = 5.77, *p* < 0.01, *BF*_10_ = 7.98) and Kalman filter learning model (*t*(614) = 5.13, *p* < 0.01, *BF*_10_ = 7.63). In previous research with a restless bandit task (Speekenbrink & Konstantinidis, 2015), the PMU decision strategy combined with a Kalman filter learning model also provided a superior fit to participants’ behaviour. Hence, the present findings could indicate that some people switched to a non-contextual strategy within this more difficult set-up.

The median parameter estimates of the GP-RBF-learning model over all acquisition functions were extracted for each participant individually and are shown in Figure 7.

**Figure 7.**
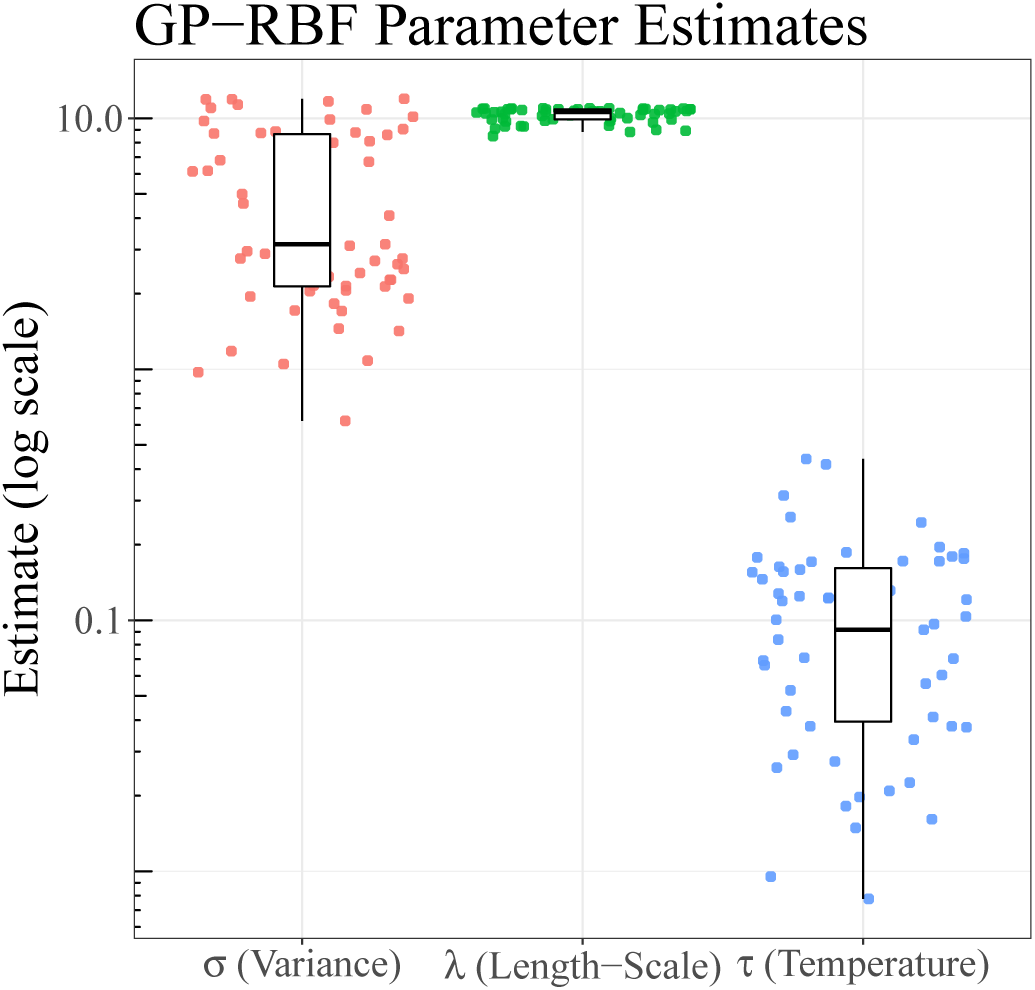
Parameter estimates of the error variance *σ*, the length-scale *λ*, and the temperature parameter *τ* for the GP-RBF model in Experiment 2. Dots show median parameter estimates per participant and boxplots show the median and inter-quartile range.

The estimated temperature parameter was 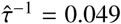 on average, which indicates that participants mostly consistently chose the options with the highest predicted utility. The estimated error variance was 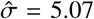 on average, which was very close to the actual variance of *σ* = 5 (*t*(58) = 0.16, *p* > 0.05, *BF*_01_ = 0.14). The estimated length-scale parameter was clustered tightly around a value of 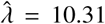. This indicates a tendency towards further extrapolation than in Experiment 1, but is still quite far removed from the level of extrapolation a linear function would provide.

## Experiment 3: Continuous-Non-Linear CMAB

The previous experiments showed that most participants were able to learn how a contexts defined by multiple features differentially affect the rewards associated to decision alternatives. The goal of the third experiment was to investigate assess whether this would still be the case in an even more complicated environment in which rewards are associated to the contexts by general non-linear functions sampled from a Gaussian process prior.

### Participants

60 participants (28 female) with a mean age of 29 (SD=8.2) were recruited via Amazon Mechanical Turk and received $0.3 as a basic reward and a performance-dependent reward of up to $0.5. The experiment took on average 12 minutes to complete on participants earned $0.67 ± 0.04 on average.

### Task and Procedure

The task was identical to that of Experiment 2, apart from the functions mapping inputs to outputs, which were drawn from a Gaussian process prior:

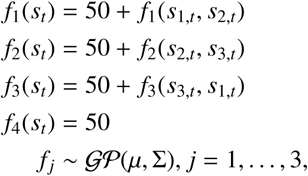

with mean function *μ* set to 0 and Σ a radial basis function kernel with a length-scale of *θ*_2_ = 2. As in Experiment 2, the features were described numerically and could take values between −10 and 10. These values were sampled from a uniform distribution 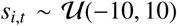. As before, the average expectation for all planets was 50 and the variance for the fourth arm was the lowest.

The procedure was identical to the one of Experiment 2.

### Results

#### Behavioural results

Participants earned 55.35 (SD =6.33) points on average during the whole task, which is significantly above chance level, *t*(59) = 5.85, *p* < 0.01. This was confirmed in a hierarchical Bayesian t-test over participants’ scores, *BF*_10_ = 54.1. 26 participants performed better than chance as assessed by a simple t-test with *α* = 0.05.

Average scores increased over trials, *r* = 0.19, *p* < 0.01, *BF*_10_ = 1.2, but to a lesser extent than in Experiment 2 (see Figure 8b), which might be due to the increase in difficulty of the task. Only 10 participants showed a significantly positive correlation between trial number and score. While significant, the increase in choosing the best option over trials was not substantial, *r* = 0.12, *p* < 0.05, *BF*_10_ = 0.3 (see Figure 8c). The proportion of choosing the non-contextual arm did not significantly decrease over time, *r* = 0.04, *p* > 0.05,*BF*_10_ = 0.1. Overall, these results seem to indicate that participants struggled more to perform well in the continuous non-linear task than in the two prior experiments.

**Figure 8.**
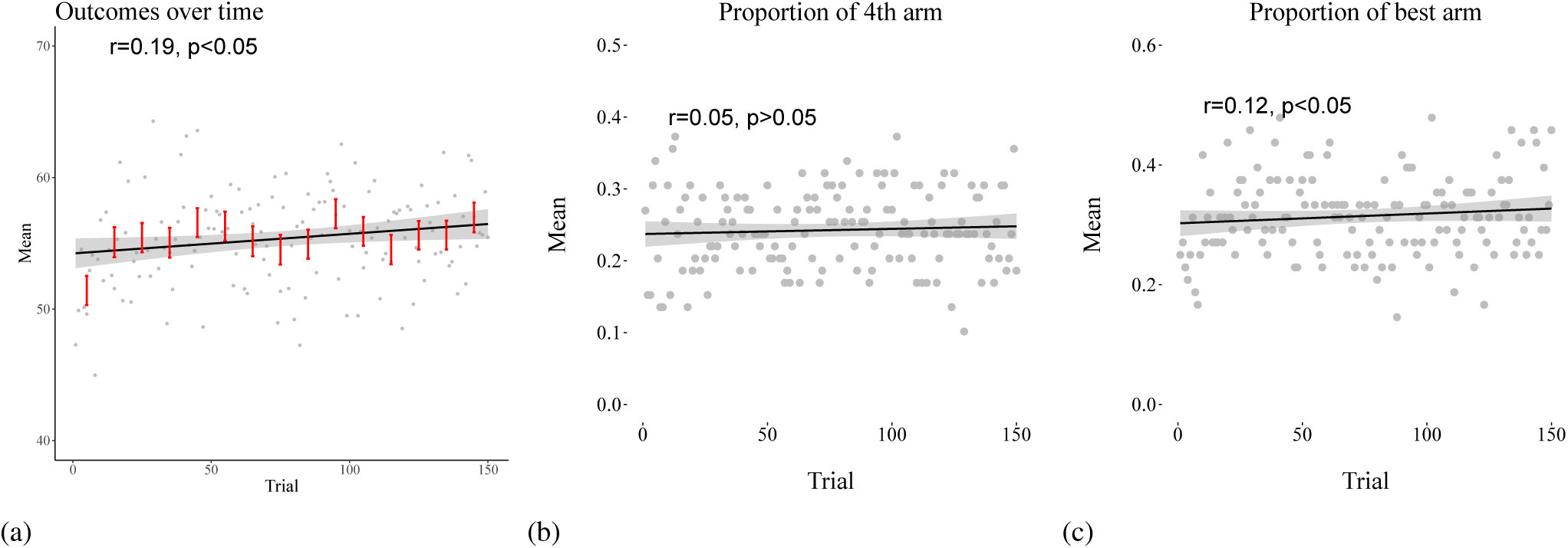
Results of the the continuous-nonlinear CMAB task of Experiment 3. (a) average score per round, (b) proportion of choices of the 4th arm, and (c) proportion of choices of the best arm. Red error bars indicate standard error aggregated over 5 trials. Regression line is based on a least square regression including a 95% confidence level interval of the prediction line.

#### Modelling results

Modelling results are shown in Figure 9. Overall, the best performing model had a GP-RBF learning component and a UCB decision strategy. Considering the results for the learning models (aggregating over the decision strategies), as in Experiment 2, the contextual models did not predict participants’ choices significantly bet-ter than the context-blind models (*t*(197) = 1.71, *p* > 0.05, *BF*_10_ = 0.13), but this was due to the linear model generating worse predictions than all the other models (*t*(197) = 3.26, *p* < 0.01, *BF*_10_ = 6.9). The GP-RBF learning model generated better predictions than the other models (*t*(197) = 3.26, *p* < 0.01, *BF*_10_ = 7.59). Regarding the decision strategy, the probability of maximum utility acquisition function generated the best predictions for both context-free models (Bayesian Mean Tracker: *t*(191) = 2.33, *p* < 0.05,*BF*_10_ = 6.87; Kalman filter: *t*(192) = 2.10, *p* < 0.05, *BF*_10_ = 7.19). The upper confidence bound sampler was the best acquisition function for the linear learning model (*t*(193) = 1.97, *p* > 0.05, *BF*_10_ = 7.53). There was no meaningful difference between different acquisition functions for the GP-RBF model.

**Figure 9.**
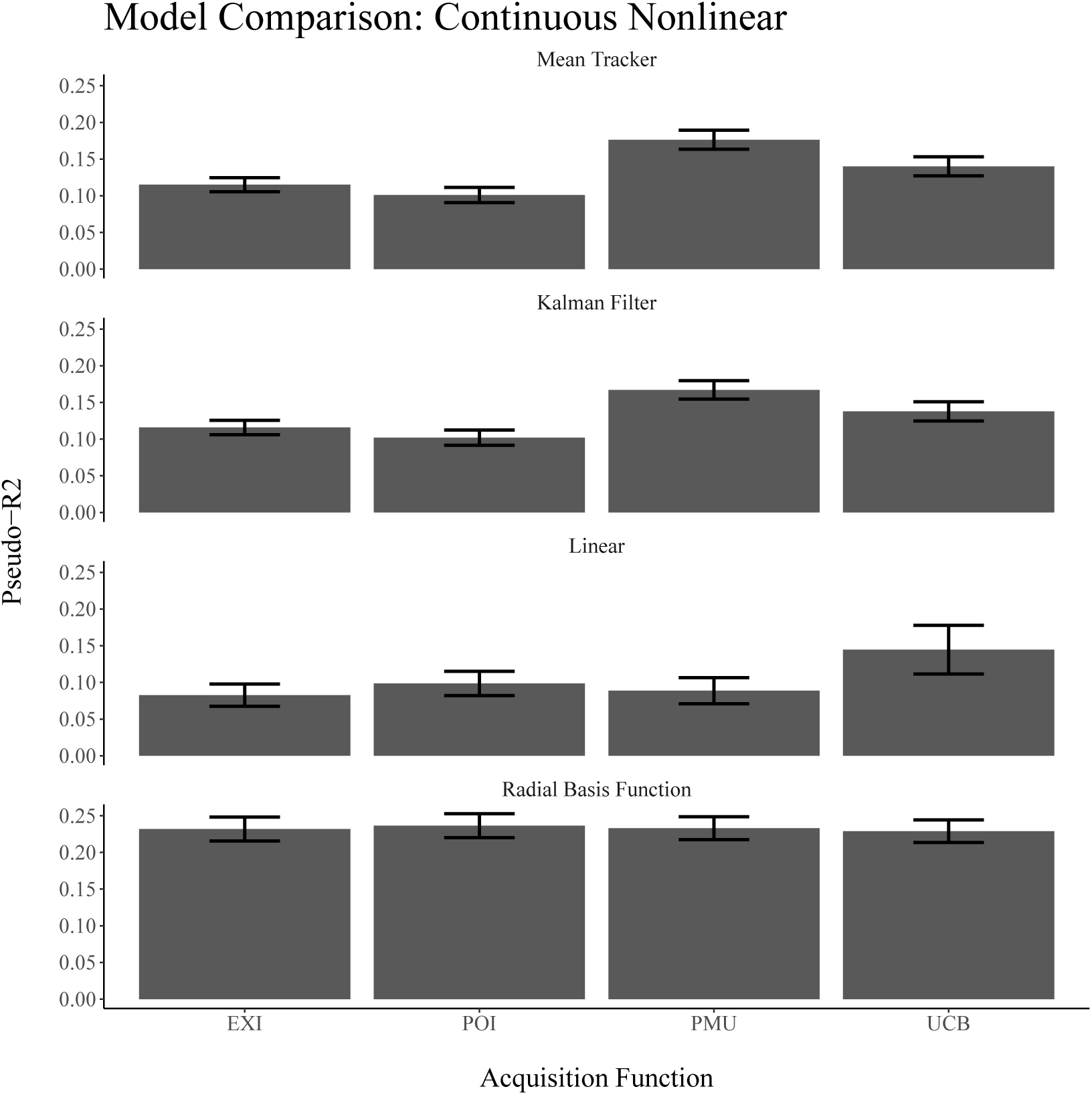
Predictive accuracy of the models for the CMAB task with continuous-non-linear cues in Experiment 3. Error bars represent the standard error of the mean.

Figure 10 shows the median parameter estimates of the GP-RBF learning model for each participant.

**Figure 10.**
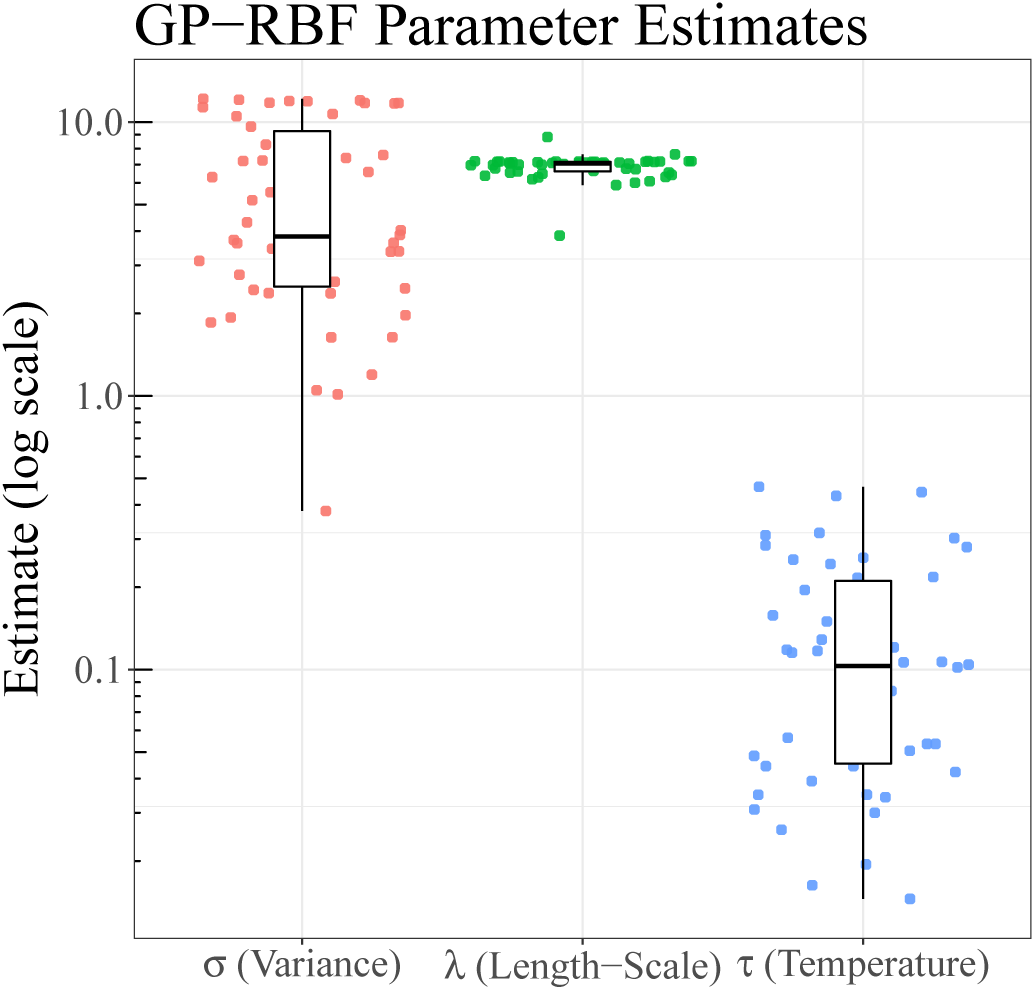
Parameter estimates of the error variance *σ*, the length-scale *λ*, and the temperature parameter *τ* for the GP-RBF model in Experiment 3. Dots show median parameter estimates per participant and boxplots show the median and inter-quartile range.

**Figure 11.**
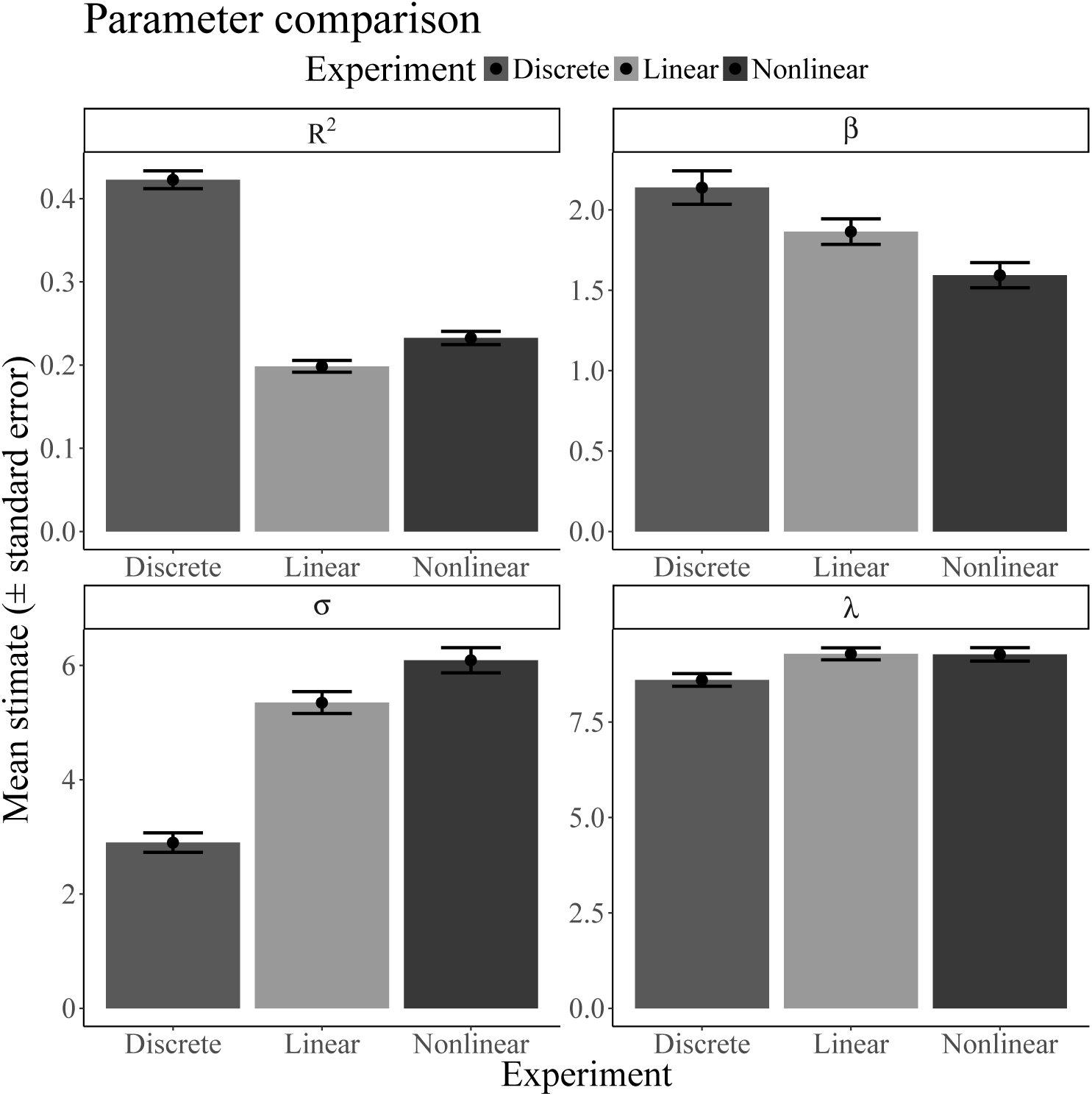
Mean estimates of the predictive performance *R*^2^, the exploration parameter *β*, the error variance *σ*, and the length-scale *λ* across all experiments. Error bars represent the standard error of the mean.

The low average estimated temperature parameter 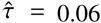 again indicates that participants mostly consistently chose the options with the highest predicted rewards. The estimated length-scale clustered tightly along a value of 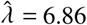, which this time turned out to be higher than the true underlying length-scale. The estimated noise variance of 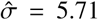 was again indistinguishable from the underlying true variance of *σ* = 5 (*t*(49) = 1.29, *p* > 0.05, *BF*_10_ = 0.34).

As this last experiment required participants to learn three different non-linear functions, it may have been too taxing for some participants to learn the functions, so that they reverted to learning in a context-free manner. Thus, whereas some participants are well-predicted by the contextual models, others seem to be captured better by the context-blind models.

## Inter-experimental model comparison

In all three experiments, the GP-RBF learning model described participants learning the best. In the first experiment, best performing model coupled this with a probability of improvement decision strategy, while in other experiment, this learning model was coupled with an upper confidence bound decision strategy. To further investigate how participants adapted to the different task environments, we here assess how model performance and parameter estimates vary between different experiments. For this analysis, we focus on the model with a GP-RBF learning component and a UCB decision strategy because this strategy described participants reasonably well in all of the experiments and come with the additional benefit that the parameters are very interpretable. For example, higher *β*-estimates are an indicator of more exploration behaviour, higher *λ*-estimates indicate further generalization, and higher noise parameters model an increasing tendency to perceive the underlying function as noisy. 11 shows the mean estimates of this model across all three experiments.

The overall predictive performance of the model was significantly higher in the first experiment compared to the other two experiments (*t*(152) = 4.52, *p* < 0.01, *BF*_10_ = 3.16). There was no meaningful difference between the continuous-linear (Experiment 2) and the continuous-non-linear tasks (Experiment 3; *t*(105) = −0.28, *p* > 0.05, *BF*_10_ = 0.24). Comparing the exploration-parameter *β* across experiments revealed that there was a negative correlation between the tendency to explore and the complexity of the task (ranked from discrete to non-linear) with *r* = −0.18, *p* < 0.05 and *BF*_10_ = 5.6. This means that participants appear to explore less as the task becomes more difficult. The assumed noise term *σ* was estimated to be lower for the discrete task than for the continuous-linear task (*t*(140) = 3.3, *p* < 0.01, *BF*_10_ = 4.35), which in turn was smaller than the estimated variance of the continuous-nonlinear task (*t*(163) = 2.22, *p* < 0.05, *BF*_10_ = 4.7). Thus, the more difficult a task, the higher the subjective level of noise seems to be. The length-scale parameter *λ* did not differ significantly between the three experiments (all *p* > 0.5, *BF*_10_ = 1.1). This indicates that participants seem to approach diverse function learning tasks with a similar assumption about the underlying smoothness of the function. While this assumed smoothness was less than the objective smoothness of of the functions in the first two experiments, it was slightly higher in the last experiment.

In summary, comparing parameter estimates of the GP-RBF model combined with Upper Confidence Bound sampling between experiments showed that (1) the model captures participants’ behaviour best for the more simple task with discrete-feature contexts, (2) participants seem to explore less in more difficult tasks, (3) the length-scale parameter which reflects the assumed smoothness of the functions seems to be relatively stable across tasks, indicating a general approach to learning about unknown functions, and (4) the continuous-non-linear experiment was hard for participants as the model captured their behaviour less well and assumed more noise overall.

## Discussion and Conclusion

We have introduced the contextual multi-armed bandit (CMAB) task as a paradigm to investigate behaviour in situations where participants have to learn functions and simulta-neously make decisions according to the predictions of those functions. The CMAB is a natural extension of both function learning and experience-based decision making in multi-armed bandit tasks. In three experiments, we assessed people’s performance in a CMAB task where a general context affected the rewards of options differently (i.e. each option had a different function relating contexts to rewards). Even though learning multiple functions simultaneously is likely to be more complex than learning a single function (as is common in previous studies on function learning and multiple cue probability learning), on average, participants were able to perform better than expected if they were unable to take the context into account. This was even the case in a rather complex situation where the functions were sampled from a general distribution of non-linear functions, although performance dropped considerably compared to simpler environments with linear functions.

Modelling function learning as Gaussian process regression allowed us to incorporate both rule-based and similarity-based learning in a single framework. In all three environments, participants appeared to learn according to Gaussian process regression with a radial basis function (RBF) kernel. This is a universal function learning engine that can approximate any functional form and assumes the function is relatively smooth. As it involves similarity-based generalization from previous observations to current contexts, it is similar to exemplar models which generalize by retrieving previously memorized instances and weighting these according to the similarity to the current context. We did not find the strong bias towards linear functions that has been found previously (e.g., Lucas et al., 2015). This could be due to the increased complexity of learning multiple functions simultaneously, or due to participants learning the functions with the purpose of making good decisions, rather than to accurately predict the outcomes as such. While good performance in standard function learning experiments requires accurate knowledge of a function over its whole domain, more course-grained knowledge usually suffices in CMAB tasks where it is enough to know which function has the maximum output for the given context. Participants appeared to assume the functions were less smooth than they actually were in the two first experiments. Although they would be expected to perform better if their assumed smoothness matched the objective smoothness, participants would have had to learn the smoothness from their observations, which is not a trivial learning problem. If the objective smoothness is unknown, approaching the task with a relatively less smooth kernel may be wise, as it will lead to smaller learning errors than overshooting and expecting relatively too smooth functions (see Schulz, Speekenbrink, Hernández-Lobato, Ghahramani, & Gershman, 2016; Sollich, 2001).

The results regarding the decision strategy were somewhat less consistent. When the features comprising the contexts were binary, people appeared to rely on a strategy in which they focus on the probability of improving upon past outcomes. In environments with continuous contextual features, they appeared to balance expectations and uncertainty more explicitly, relying on an upper confidence bound (UCB) acquisition function. Participants may have adapted their decision strategy to the task at hand. In a relatively simple scenario with binary features and small number of unique and distinct contexts, it is feasible to memorize the average rewards and best alternative for each context, and trying to maximally improve upon the current best option may therefore be an efficient strategy. As the environment becomes more complicated, memorization seems less plausible, making exploration in order to learn the functions more important. The UCB strategy explicitly balances the expected rewards and its associated uncertainty, and has been interpreted as a dynamic shaping bonus within the exploratory choice literature (Daw, O’Doherty, Dayan, Seymour, & Dolan, 2006). It is currently the only acquisition function with provable good regret (Srinivas, Krause, Kakade, & Seeger, 2012).

The environment involving non-linear functions sampled from a Gaussian process was more difficult than the others, and a proportion of participants appeared unable to learn the functions. Their behaviour was more in line with a context-blind learning strategy (Kalman filter) that treats the task as a restless bandit in which the expected rewards fluctuate over time but where these fluctuations are not predictable from changes in context. The combination of a Kalman filter learning model with a “probability of maximum utility” decision strategy that described these participants best has been found to describe participants behaviour well in an actual restless bandit task Speekenbrink and Konstantinidis (2015) and here might have indicated the limits of participants’ learning ability in our task.

The present experiments focused on a general context which differentially affected the outcomes of options. This is different than the CMAB task of Stojic et al. (2015), in which the features had different values for each option, while the function relating the contexts to rewards was the same for each options. Future studies could combine these paradigms and incorporate both option-specific (e.g., the type of restaurant) as well as general (e.g., the area in which the restaurants are located) contextual features, possibly allowing these to interact (e.g., a seafood restaurant might be preferable to a pizzeria in a fishing village, but not a mountain village).

To make bring our task closer to to real-life decision situations, future research could adapt the reward functions to incorporate costs of taking actions or obtaining poor outcomes (see Schulz, Huys, Bach, Speekenbrink, & Krause, 2016). Research utilizing the CMAB paradigm also has the potential to be applied to more practical settings, for example military decision making, clinical decision making, or financial investment scenarios, to name just a few examples of decision making that normally involve both learning a function and making decisions based on expected outcomes. Incorporating context into models of reinforcement learning and decision making generally provides a fruitful avenue for future research.

## Footnotes

1 Normally, the algorithm would pick the observation with the highest value according to the acquisition function, whereas we enter these values into a softmax function, see Equation 17

2 In earlier studies (Schulz, Konstantinidis, & Speekenbrink, 2015) we had implemented Thompson sampling as sampling functions from the Gaussian process and individually maximizing the resulting functions instead of sampling from the posterior predictive distribution. We also did not estimate hyper-parameters for the Gaussian process for each participant.

3 The initial trial had the same context *s*_1_ for all participants. Afterwards, the values of the context *s_t_* were sampled at random

4 https://github.com/ericschulz/contextualbandits

5 As previous research with the Iowa Gambling task found little effect of options’ position on participants decisions (Chiu & Lin, 2007), we expect similar results if we had randomized the position on screen.

6 Implemented as a Bayesian meta t-test that first compares each participant’s scores against 50 and then aggregates the overall results in a Bayesian meta t-test (see Morey et al., 2015).

